# Designing the database for microarray experiments metadata

**DOI:** 10.1101/101956

**Authors:** Oleksandr Lykhenko, Alina Frolova, Maria Obolenska

## Abstract

Advancements in both computer science and biotechnology opened way to unprecedented amount and variety of gene expression studies raw data in the open access. It is sometimes worth to rearrange and unite data from several similar gene expression studies into new case-control groups to test new hypothesis using available data. Unfortunately, most popular gene expression databases such as GEO and ArrayExpress were not designed to allow such cross-study procedures. In order to locate comparable samples in different studies numerous steps are required including gathering additional sample metadata and its standardization. Specialized databases are developed by investigators in their own fields of interest to reuse the processed data and create different case-control groups and test multiple hypothesis.

Here we present detailed description of the specialized database creation along with its use case which is 32 gene expression cDNA microarray datasets on human placenta under conditions of pre-eclampsia containing expression data on more than 1000 biological samples. Samples contain sufficient metadata for them to be merged into relevant cross-experiment case-control groups for further integrative analysis.

## Background

Current database is being developed with intention to gather clinical, biological and other metadata for biological samples of the microarray experiments on preeclampsia-affected human placenta available at ArrayExpress. Main purpose of gathering these metadata is the construction of cross-experiment case-control sample groups for subsequent integrative analysis. Despite the fact that ArrayExpress stores the considered data already, there are several reasons to maintain a database separate from ArrayExpress. First, names and values in samples’ metadata are not standardized, since they were uploaded independently and at different times. For example, the fetal gestation age can be referred as age, fetal age, fetus age (wks) and in other similar ways, and therefore gestation age for all samples in a set of experiments can not be obtained via simple search query. Second, metadata is often incomplete and insufficient for the samples to be comparable in terms of into which case or control group they should be placed. Third, ArrayExpress will most certainly not allow third-party metadata update requests, let alone our suggestions on metadata standardization.

Although our database can be used to store data on any cDNA microarray experiments or, indeed, any experiment that uses technical means to process some amount of samples, the database was designed for specific task, which is an integrative analysis of microarray experiment datasets on preeclampsia-affected human placenta.

Pre-eclampsia [1] is a disorder that occurs only during pregnancy and the post-partum period and affects both the mother and the unborn baby. Affecting at least 5-8% of all pregnancies, it is a rapidly progressive condition characterized by high blood pressure and the presence of protein in the urine. Typically, pre-eclampsia occurs after 20 weeks gestation (in the late 2nd or 3rd trimesters or middle to late pregnancy) and up to six weeks postpartum (after delivery), though in rare cases it can occur earlier than 20 weeks. Globally, pre-eclampsia and other hypertensive disorders of pregnancy are a leading cause of maternal and infant illness and death.

Although etiology and pathogenesis of pre-eclampsia are still unknown numerous studies (listed in Supplement 1) point at the gene disregulation in placenta to be the cause of the disease. Besides, there are also known cases when placenta develops without fetus, called molar pregnancy [5], which are associated with very early-onset pre-eclampsia. This is why we focused our study on gene expression in placenta. Finally, we chose cDNA microarray technology as most popular in pre-eclampsia studies among ones providing information on the whole transcriptome.

## Methods

Our software is written in Python language using Django framework for web interface development and Postgres for relational database support. All experiment and sample metadata were automatically extracted from ArrayExpress database via Bioservices which is a Python interface to ArrayExpress. NCBI database was used to supplement the missing data along with the corresponding scientific articles and authors personally.

Source code and database backup file are available at GitHub: https://github.com/Sashkow/placenta-preeclampsia Web interface for our database at its current stage can be accessed at: http://194.44.31.241:24173/

## Database schema development

As we are able to download the microarray experiment description, the microarray platform description and a table of sample-wise metadata for each experiment from ArrayExpress, we have the following entities/tables: **Experiment, Microarray, Sample.** The first naive approach for the database schema is in Figure 1. The catch here is three dots in each entity. While Experiment and Microarray entities tend to have the characteristics uniform across ArrayExpress, the Sample entities are likely to have different set of attributes depending on Experiment they belong to. Hence, each new sample characteristic (here and later, attribute) will require database schema update.

**Figure 1:**
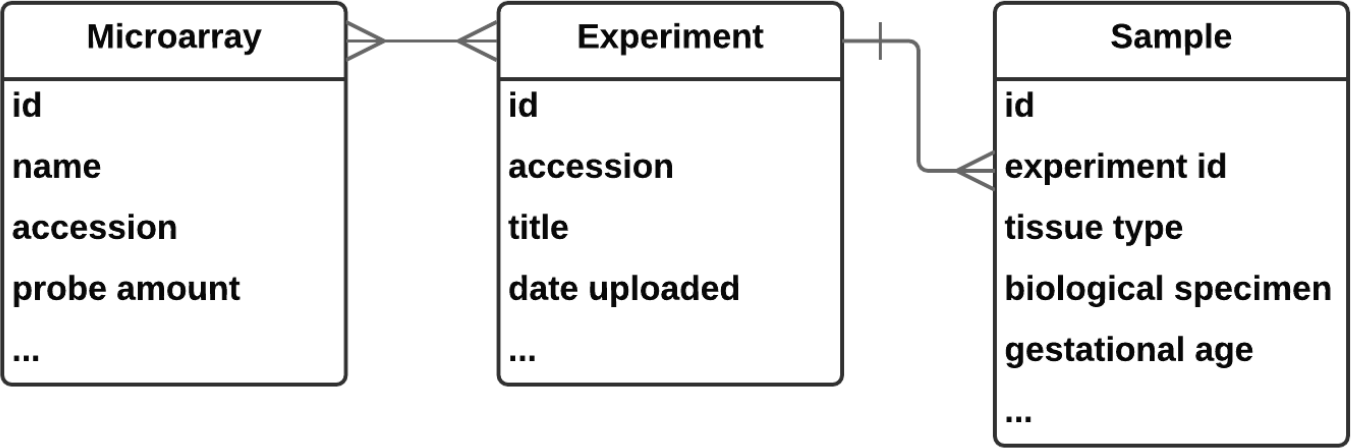
Initial database entity relation diagram

One way to address this issue would be the choice of schemaless document oriented database like MongoDB. Unfortunately, our selection of web development tools, which are Python 3 with Django framework, inclined us to choose Postgres as more compatible with Python/Django.

Luckily, Postgres itself provides us with plenty of schemaless approaches [10, 13]. Postgres has HStore custom field which is essentially a dictionary with strings for keys and values. This updated our database to be practically schemaless as can be seen in Figure 2.

**Figure 2:**
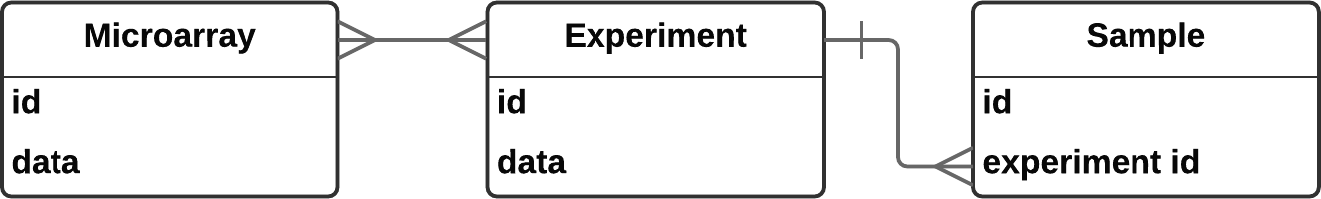
Almost schemaless database; ”data” is an HStore field.

This way if we have a sample with the attribute ”age” of value ”37 weeks” and attribute ”diagnosis” of value ”pre-eclampsia” downloaded from ArrayExpress we store it in Samples table with the ”data” field equal to ””age”:”37 weeks”, ”diagnosis”:”pre-eclampsia””.

However, this implementation turned out to have drawbacks. When we uploaded our data and began sample attribute names and values standardization it occurred to us that we want to store the originally downloaded names and values along with the standardized ones. To this end sample attributes can no longer be string name-value pairs in Sample’s *data* HStore dictionary field, as we now want to store four strings instead of two for each sample attribute: old name, old value, new name, new value. And, since Django does not allow nested HStore fields, *data* field should be replaced with separate entity named SampleAttribute containing SampleAttributeName and SampleAttributeValue (Figure 3)

**Figure 3:**
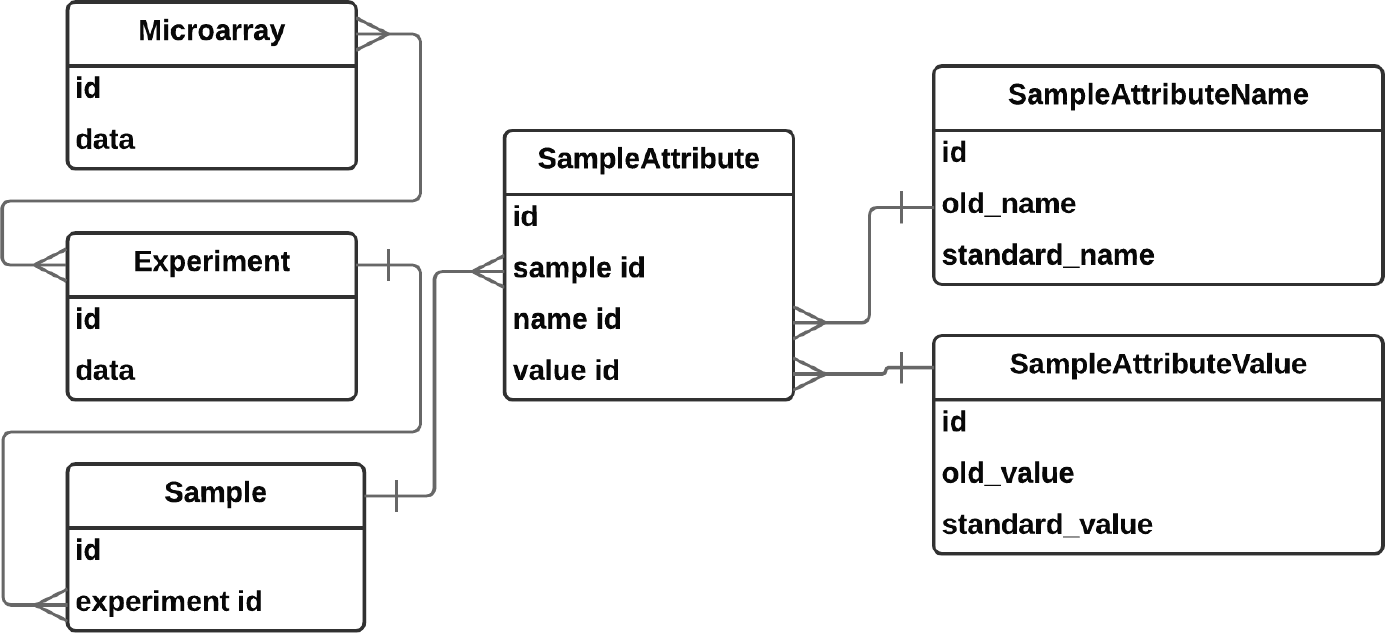
Schema modified for keeping original values

This schema leads to massive duplication of data because the ”original-standard” mapping is ambiguous meaning that the same original name may map onto multiple standard name depending on the context of the experiment. For example, original name ”placenta” may map to ”Chorionic Villi” standard name if the corresponding article mentions this exact tissue type or just to ”Placenta” otherwise.

In this situation it may be a good idea for standard names and values to be described once and then referred everywhere else throughout database. Old names and values, on the other hand, should be sample-attribute-related. The following schema (Figure 4) implements that.

**Figure 4:**
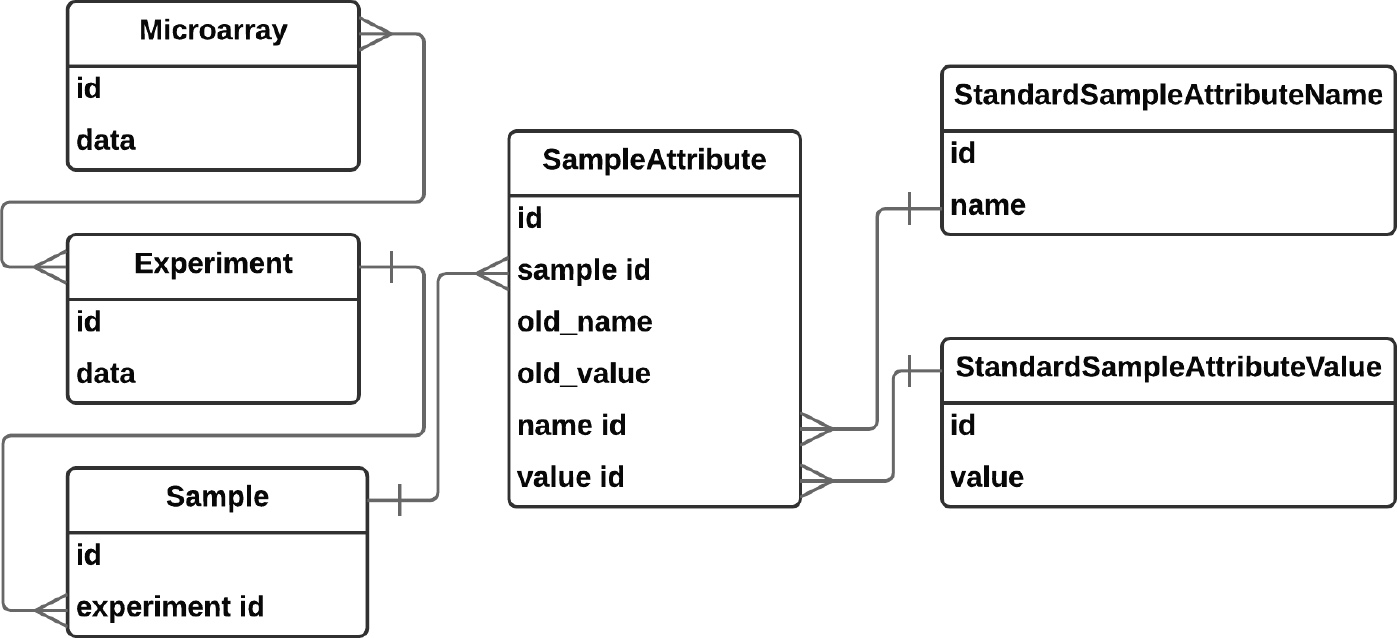
Schema modified to avoid data duplication

During the sample attribute standardization procedure we found out that not all original names and values can be mapped to a MeSH term as we intended. We then began to use other ontologies such as Experimental Factor Ontology (EFO) and to define our own terms. To satisfy our need in storing additional info about StandardSampleAttributeName and StandardSampleAttributeValue we added *additional_info* HStore field to these entities. To reflect the fact that some standard names and values are related either as *synonyms* or as parent-child we added synonyms reflexive (to oneself) many-to-many field to StandardSampleAttributeName and StandardSampleAttributeValue entities. Finally, to show the relation between standard values and names we added one-to-many relationship from StandardSampleAttributeValue to StandardSampleAttributeName. The results are in Figure 5.

**Figure 5:**
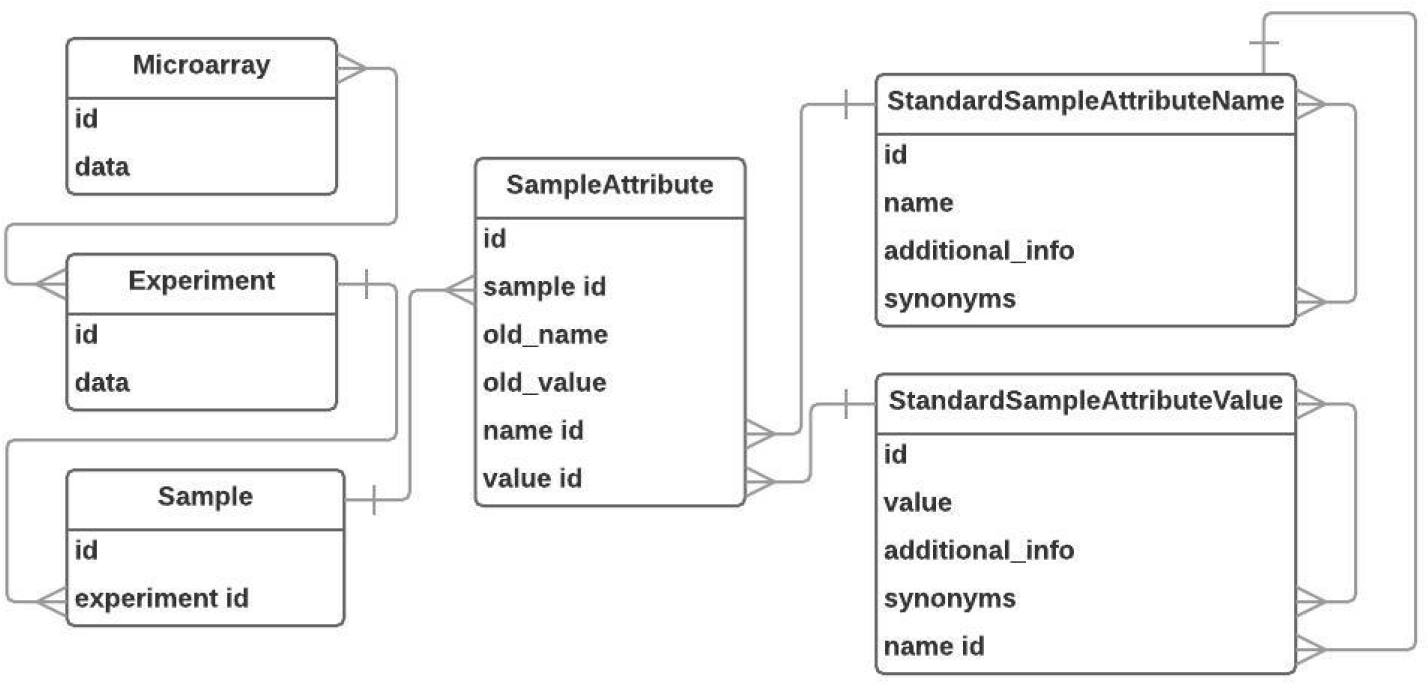
Final Schema

### Important note

Not all sample attribute values are qualitative with restricted list of possible values. Some of them are quantitative with numeric, textual or Boolean values. For quantitative values actual value is stored in *old value* field while the standard value specifies value type and units (like ”number in weeks”, ”mass in grams”). Thus, a Gestation Age sample attribute may have standard value of Number in weeks and and an old value of 42.

### Another important note

Some attributes’ original names and values were excluded from standardization process and mapped to a special standard name (or value, respectively) named ”(excluded)” either due to irrelevance of the information in those attributes or because that information was moved to other attributes. Those were: ”Unknown Sex”, ”Other”, ”cal id”, ”passage”, ”matching”, ”N/A” ect.

Originally blank names and values were filled with ”<empty>” value.

## Use case: filling database with microarray experiments on preeclampsia-affected placenta

The initial list of 43 relevant datasets was obtained as a result of ArrayExpress search by the following query: ”preeclampsia OR pre-eclampsia OR preeclamptic OR pre-eclamptic” with results filtered by organism ”Homo sapiens”, experiment type ”rna assay”, experiment type ”array assay”. E-GEOD-25906 was excluded due to data retrieval failure. E-MTAB-3732 was excluded since it is a compilation of microarray experiments for different diseases taken from publicly available sources. E-GEOD-15787, E-GEOD-22526, E-MEXP-1050 were excluded due to old microarray design or failure to find probe nucleodide sequences for the array. The full list of included and excluded can be found in Supplement 1.

From the chosen 32 (more than 1000 samples) datasets we downloaded relevant metadata and thus automatically filled Experiment, Microarray, Sample and original name-values for SampleAttribute entity. We then manually mapped original names and values to the standard ones according to well known ontologies such as MeSH and EFO [3, 6]. Missing metadata was mined from the text of the corresponding articles and from authors personally.

Most of the considered samples are comparable and ready for integrative analysis at this moment. These biological samples can now constitute case-control groups of larger size than in original datasets.

## Discussion

Numerous attempts to organize gene expression data have been taken in due time including the databases we took our data from: ArrayExpress and GEO.

**G**ene **E**xpression **O**mnibus (**GEO**), an NCBI’s gene expression repository founded in 2000, has a straight forward general data structure [11].

**Figure 6:**
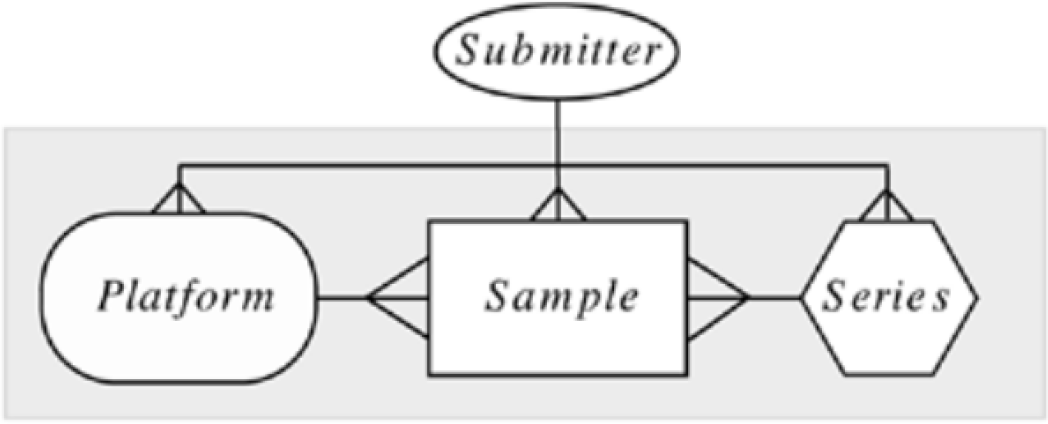
GEO entity relation diagram

*Platform* entity contains a list of probes that define what set of molecules may be detected in any experiment utilizing that platform. An instance of *sample* describes the derivation of the set of molecules that are being probed and utilizes platforms to generate molecular abundance data. Each sample has one, and only one, parent platform which must be previously defined. Sample may include submitter-defined clinical and other metadata concerning the sample although no standard sample attribute naming format is imposed onto submitter. *Series* entity unites samples of the same experiment. It usually includes experiment related metadata such as reference to a corresponding article.

A need for a more well-annotated experiment and sample data as well as necessity to embed external data from numerous specialized databases in accordance with already established community standards led to creation of **ArrayExpress** gene expression database in 2002. [14]. Unlike GEO, database structure of ArrayExpress includes over 200 unique tables for various input data formats. Web-access on the contrary is simple and the entities accessible to user are similar to ones in GEO: experiments, samples, microarray platforms, protocols. Means of programmatic data access (also known as ”application programming interface” or just API) are also available for ArrayExpress [7]. API allows retrieving detailed metadata associated with an experiment: sample annotation, protocols and data files - in XML format without need for looking through multiple tables directly.

ArrayExpress provides ability to search for specific sets of experiments depending on experiment type, submission date, studied organism, experiment description. Search query keywords are looked up in Experimental Factor Ontology to extend search with synonyms and alternative forms of the keywords (not only ”pre-eclampsia”, but also ”preeclampsia”, ”pre-eclamptic”).

A more recent **BioSamples** database has been developed focused on search for samples with similar characteristics across different experiments [2]. This database, however has its own limitations due to the absence of standard sample attribute naming and to incomplete annotation, which is an obstacle to constructing cross-experiment study groups.

**GENEVESTIGATOR** is a search engine for gene expression over a compendium of manually curated datasets [4, 12, 16]. Biological samples are carefully annotated using custom ontology with a variety of characteristics allowing to filter for highly specific cases. Cross-experiment data search and analysis is based on a concept of meta-profile. As explained in [9] meta-profiles summarize expression levels from many samples according to their biological context. Each sample is annotated with five attributes: anatomical part, cell line, cancer type, developmental stage, and perturbation - to generate meta-profiles. An exception is the Perturbation meta-profile, which consists of responses to various experimental conditions (drugs, chemicals, hormones, etc.), diseases, and genotypes. For Perturbation meta-profile results are created by comparing groups of samples from individual experiments. Data from multiple experiments are not mixed to create a single value. As a result, this tool contains large compendia of response types collected from many experiments. It also worth mentioning that samples of two different microarray platforms can not be considered simultaneously.

Another resource enabling interactive query and navigation of transcriptome datasets is **Gene Expression Browser** [15] and its specific implementation for placental gene expression [8]. While its data is more relevant to our pursues and the interface is quite convenient and has some extended features in comparison to GEO or ArrayExpress, such as search for differentially expressed genes in a single experiment, building relevant plots, giving detailed information on found genes and and a bit of meta-analysis tools such as search for experiments that have given gene differentially expressed with a certain rate of difference, no tools for integrative analysis are provided.

At this time most of the samples in our database are provided with sufficient metadata for them to be comparable. These biological samples contain more complete and standardized metadata than ArrayExpress can offer and thus the samples can now constitute case-control groups of larger size than in individual datasets. Described here database shall be used in forthcoming analysis, namely, the search for differentially expressed genes between different cross-experiment case and control groups. Also, our database is designed and expected to be enlarged with forthcoming gene expression experiments’ metadata and, potentially, with data from other omics (genome, metabolome) with relatively low effort.

## Author’s contributions

OL, AF designed the database. AF, MO made suggestions on database content. OL implemented database design and its admin web interface and performed metadata standardization. All authors read and approved the final version of the article.

## Additional Files

Supplement 1 – ArrayExpress Experiments’ Accession Numbers

